# Latent functional connectivity underlying multiple brain states

**DOI:** 10.1101/2021.04.05.438534

**Authors:** Ethan M. McCormick, Katelyn L. Arnemann, Takuya Ito, Stephen José Hanson, Michael W. Cole

## Abstract

Functional connectivity (FC) studies have predominantly focused on resting state, where ongoing dynamics are thought to reflect the brain’s intrinsic network architecture, which is thought to be broadly relevant because it persists across brain states (i.e., is state-general). However, it is unknown whether resting state is the optimal state for measuring intrinsic FC. We propose that latent FC, reflecting shared connectivity patterns across many brain states, better captures state-general intrinsic FC relative to measures derived from resting state alone. We estimated latent FC independently for each connection using leave-one-task-out factor analysis in 7 highly distinct task states (24 conditions) and resting state using fMRI data from the Human Connectome Project. Compared to resting-state connectivity, latent FC improves generalization to held-out brain states, better explaining patterns of connectivity and task-evoked activation. We also found that latent connectivity improved prediction of behavior outside the scanner, indexed by the general intelligence factor (*g*). Our results suggest that FC patterns shared across many brain states, rather than just resting state, better reflects state-general connectivity. This affirms the notion of “intrinsic” brain network architecture as a set of connectivity properties persistent across brain states, providing an updated conceptual and mathematical framework of intrinsic connectivity as a latent factor.

## Introduction

A major goal in cognitive neuroscience in recent years has been to move away from characterizing brain activation and connectivity in specific task states towards understanding “intrinsic” or context-free brain activity. Such activity reflects the more than 95% of metabolic brain activity that remains unchanged across cognitive demands (Raichle, 2006). This ongoing brain activity persists across states and is not attributable to external stimuli or task demands. Efforts to understand intrinsic function have focused primarily on statistical associations between brain activity time series (functional connectivity; FC) during the resting state (Fox & Raichle, 2007) (but see (Finn et al., 2015; Greene, Gao, Scheinost, & Constable, 2018) for task-based investigations), which has revealed an intrinsic brain functional network architecture that recapitulates patterns of task-evoked brain activity (Cole, Ito, Bassett, & Schultz, 2016; Smith et al., 2009) and structural connectivity (Honey et al., 2009). As the name implies however, resting state is just one state that the brain can occupy, and a truly “intrinsic” connectivity network should persist across the many different states a brain might assume. In other words, a “state-general” intrinsic network. Despite its importance for understanding brain function, many uncertainties remain on how to best estimate intrinsic FC. While some efforts have focused on the need to obtain longer resting-state scans (Anderson, Ferguson, Lopez-Larson, & Yurgelun-Todd, 2011; Elliott et al., 2019; Hacker et al., 2013; Laumann et al., 2015) more recent approaches have highlighted advantages of combining resting-state and task data to analyze intrinsic activity.

This second set of approaches leverages functional data across different task (and rest) scans in order to improve the reliability of FC estimates and their predictive utility (Elliott et al., 2019) Because of the relatively high stability of FC networks across task states (Cole, Bassett, Power, Braver, & Petersen, 2014; Gratton et al., 2018; Krienen, Yeo, & Buckner, 2014), combining data across task runs aims to distinguish what is common across a larger set of brain states. What is common therefore reflects the intrinsic patterns of covariance in the brain, while variation between different brain states is treated as noise in the combined data. However, this work largely relies on averaging data from multiple scans together (Elliott et al., 2019). While this approach has been shown to be useful, and has the advantage of simplicity, there are potential theoretical limitations to such an approach that may limit its generalizability. Given its ubiquity and close-formed, arithmetic solution, the average is rarely thought of as a formal statistical model. However, recent work (McNeish & Wolf, 2020) has shown that the average can be thought of as a restricted case of the more-general factor analytic model. Embedding the average in a theoretically rich statistical framework is likely to offer advantages for interpretation of results using this measure as well as insights into the measure itself.

Factor analysis has a long tradition in the behavioral sciences (Spearman, 1904; Thurstone, 1935) and is an invaluable tool in psychometrics and psychological measurement. Its key insight is that observed measures (e.g., behavioral responses or fMRI scans) are imperfect manifestations of an unobserved (i.e., latent) variable (Bollen, 2002). In the factor model, observed indicators (***y_i,t_***; *i* = individual, *t* = task state) are modeled as dependent on the underlying latent factor (***η***; Figure 1). Variability in the indicators is partitioned into common variance (transmitted through the factor loading matrix, ***Λ***) and unique variance (***ɛ_t_***). In this model, latent FC represents an unmeasured, underlying brain state that is common to all observed brain states (i.e., the indicators: resting state, motor task, etc.), but we also explicitly model additional variance that is only found in each individual task state through the error terms. Factor loadings for the individual task states (e.g., *λ*_11_ for Rest) in this single-factor model can be interpreted as the proportion of variance explained in each task state by latent FC (similar to *R^2^* in regression).

**Figure 1.**
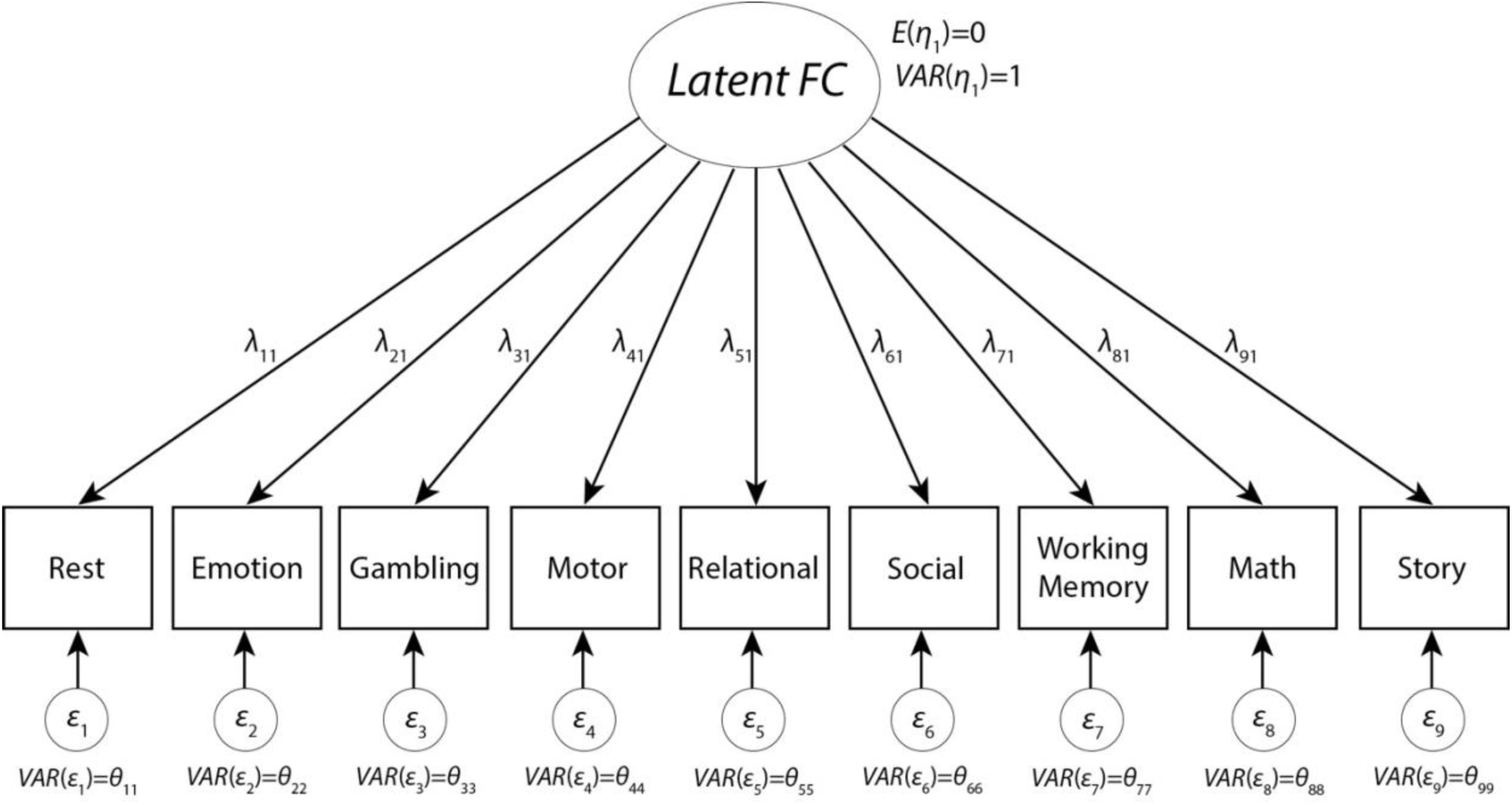
Factor Model. A set of indicators (e.g., Rest, the Motor task, etc.) are modeled being composed of shared underlying variance, as represented by the latent factor (i.e., Latent FC), and unique task-state variance (in the errors). Factor loadings (λ) represent the percent variance in each task state that is explained by the underlying Latent FC.

As can be seen in Figure 1, the factor analysis model of latent FC is a parameter-rich model that allows for differentially weighted relationships between the underlying latent connectivity and measured connectivity in each specific state. What McNeish and Wolf (2020) showed, however, is that the average can be recovered using this model by setting all factor loadings (λ) equal to 1 and the unique variances to 0. This recast of the average as a special case of the factor model not only has the advantage of making the assumptions of the average clearer, but it enables a formal test of those assumptions. For instance, by setting all factor loadings equal, the average assumes that each observed FC state is equally (and positively) related to the underlying latent FC. If we want to relax that assumption, the factor analytic model can be used to compute unique optimally weighted values for each factor loading, which suggests that some observed states may be better (or worse) reflections of underlying latent FC. Indeed, factor loadings may take on negative values, which implies that an observed indicator is anti-correlated with the underlying latent FC. However, if the assumption of equal, positive weighting is indeed an appropriate assumption, freely estimated factor loadings will converge towards equal values and approximate the average. In other words, the flexibility of the full factor loading does not preclude the average, but instead offers a broader range of possibility for deriving a measure of latent FC in heterogeneous data and can be used to test the validity of the average FC assumption of equal positive factor loadings across brain states.

Here, we test the reliability of a factor analytic framework for modeling state-general brain connectivity – “intrinsic” FC that generalizes across a variety of brain states. First, we hypothesized that latent FC reflects a positive manifold (analogous to the positive correlations across intelligence tests in general intelligence research; (Kovacs & Conway, 2016)), where all state-specific connectivity values are positively correlated with each other and so load positively onto the underlying latent variable. This would confirm that a single common intrinsic functional network architecture exists across conscious brain states. Importantly, this differs from the idea that states are correlated (Finn et al., 2015; Gratton et al., 2018) as between-subject variance is decomposed at each individual connection rather than correlating across connections. We further hypothesized that by combining information across task states, such as in the factor model, a more reliable measure of “intrinsic” connectivity can be estimated than when using resting state data alone (the current field standard). This would suggest that resting-state FC is not necessarily the best state for estimating intrinsic FC, especially if resting state does not load higher on the latent variable than other states. In testing these hypotheses, we developed an analytic framework for estimating state-general, latent FC in whole-brain functional data. Using multi-task fMRI data from the Human Connectome Project (HCP), we compare the ability of latent and resting-state FC to predict task-evoked activation and task-state FC for held-out brain states, as well as to explain individual differences in psychometric “g” (a measure of human intelligence derived with a similar factor analytic model). Results demonstrate the promise of the latent variable approach in functional neuroimaging, particularly for the estimation of intrinsic FC that generalizes beyond specific brain states (e.g., rest). Finally, we demonstrate the relationship between freely estimated latent FC and the simpler average FC approach and discuss the theoretical advantages of casting both methods in the latent variable framework for future work.

## Results

### Factor analysis model of latent connectivity

We ran independent factor analysis models for each connection, estimating the factor loadings of the latent variable (i.e., latent FC) onto each state. Latent FC captures the shared variance in FC across all states (see Figure 2). Factor analyses were run using all available data (i.e., the full time series and all states). All analyses were performed in the exploratory sample independently and then replicated in the validation sample (both N = 176; see the Participants section for additional information). Importantly, all factor analytic models were fit for each sample separately to avoid issues of circularity when comparing results across samples.

**Figure 2.**
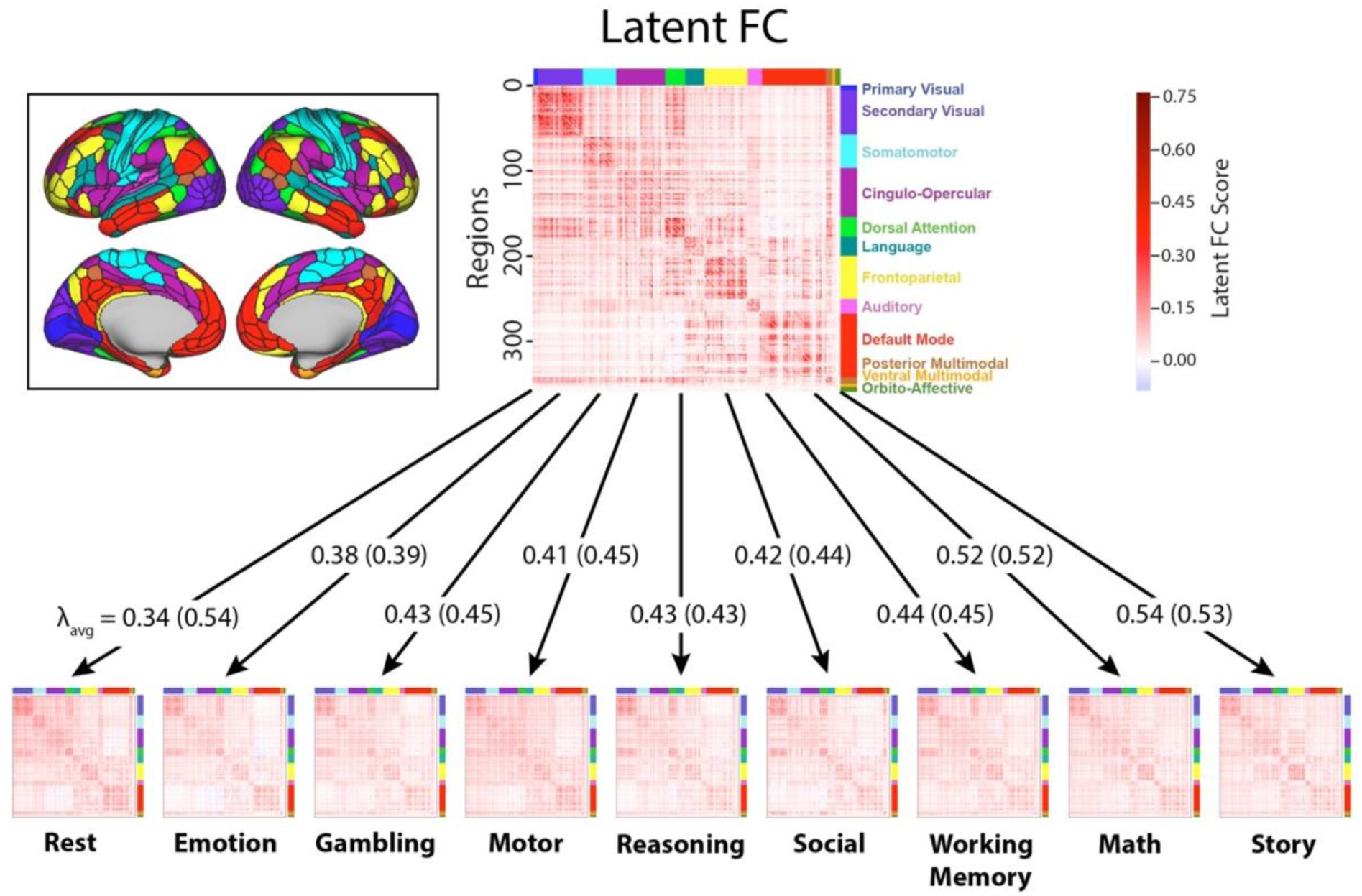
Factor analysis model of latent FC. Visualization of the latent connectivity matrix and state-specific functional connectivity matrices (group average across subjects). Color along the axes of each matrix corresponds to the network membership of each ROI. For each arrow, the average loadings (λ_avg_) for each state are shown for analyses controlling for number of TRs (first) and when not controlling for TRs (in parentheses). The averaging loadings for the task states were largely stable across analyses, but the average loading for resting-state increased substantially (from 0.34 to 0.54) when not controlling for the number of TRs. The amount of resting-state data per participant went from 4800 TRs (58 minutes) to 2112 TRs (25 minutes) when matching the total amount of “on-task” time. The network mapping is shown in the cutout (left). Elements in the state-specific matrices represent correlations (r) between regional time series and elements in the latent FC matrix represent factor scores computed from the model for each connection.

Consistent with our hypothesis that there is a “positive manifold” demonstrating a common latent FC architecture across states, almost all factor loadings were positive (greater than 99%) across all connections and all states (see Table 1). Furthermore, 70.7% of all factor loadings were reasonably large in magnitude (factor loading ≥ 0.4) and 97.4% of connections had two or more states with factor loadings ≥ 0.4 in the full latent FC model. The emotion task had the fewest large factor loadings (47.3%) and the resting state had the most (92.6%) (see Table 1 for full details).

**Table 1:** Factor Loadings. Almost all factor loadings were positive regardless of whether all resting state data were used (left) or we controlled for the number of time points between task and rest (right). Only resting state showed a substantial shift in the percent of factor loadings ≥ 0.4 when controlling for the number of timepoints. The amount of resting-state data per participant went from 4800 TRs (58 minutes) to 2112 TRs (25 minutes) when matching the total amount of “on-task” time.

|  | **All Data** |  | **Controlling for # Timepoints** |  |
| --- | --- | --- | --- | --- |
| **State** | **% Loadings $\geq 0$** | **% Loadings $\geq 0.4$** | **% Loadings $\geq 0$** | **% Loadings $\geq 0.4$** |
| Rest | 99.9% | 92.6% | 99.0% | 31.6% |
| Emotion | 99.3% | 47.3% | 98.7% | 46.3% |
| Gambling | 99.6% | 65.0% | 99.1% | 62.2% |
| Motor | 99.8% | 68.0% | 99.4% | 54.8% |
| Reasoning | 99.5% | 62.1% | 99.1% | 62.3% |
| Social | 99.8% | 66.2% | 99.3% | 58.4% |
| Working Memory | 99.7% | 67.0% | 99.2% | 64.9% |
| Math | 99.9% | 82.4% | 99.6% | 82.4% |
| Language | 99.9% | 86.0% | 99.7% | 86.3% |

To control for differences between states in the amount of data used to obtain state-specific FC estimates, factor analyses were re-run while matching the number of time points from rest and task data (2112 TRs from rest and 264 TRs for each of the 8 tasks). With this approach, resting state had the fewest number of relatively high magnitude factor loadings of all states – only 31.6% of resting state connections had factor loadings ≥ 0.4. Thus, resting state had the highest factor loadings onto latent FC when a large amount of data was used to estimate resting-state FC, but the lowest factor loadings when less data was used. Controlling for the number of time points between task and rest led to less pronounced changes in the factor loadings of the other states (see Figure 2), likely because there was no relationship between the number of TRs for a given task state and its average factor loading in the full TR analysis (see Figure S1). Note that this drop occurs even though rest continues to have substantially more TRs (8x) than any given task state in these analyses.

### Latent FC improves generalization to connectivity of held-out states

We next sought to test our second hypothesis: A more reliable and generalizable measure of “intrinsic” connectivity can be estimated by combining information across task states, such as in the factor model, than by using resting state data alone (the current field standard). To test whether the measures of intrinsic FC persist across brain states, we quantified the generalizability of rest FC and latent FC to held-out brain states. To calculate the similarity of FC patterns (i.e., across 64,620 network connections), we computed the Pearson’s correlation of rest FC or latent FC with state FC for each individual subject, applying Bonferroni correction to correct for multiple comparisons. For latent FC, similarity was always computed for the state that was held-out while running the factor analysis model. Compared to rest FC, we found that latent FC exhibited significantly greater similarity with a variety of independent brain states (see Figure 3A). Similarity of each state with latent FC was comparable across states, exhibiting the greatest similarity to the WM task (*r* = 0.71) and the least similarity to the social task (*r* = 0.66) and resting state (*r* = 0.65). Rest FC exhibited the greatest similarity to the full resting state data (*r* = 0.73), providing a measure of test-retest similarity of rest FC (i.e., how well the restricted TR data represents the correlation matrix computed on the complete resting state data). For the task states, rest FC had the greatest similarity to the motor task (*r* = 0.61) and the least similarity to the relational task (*r* = 0.56).

**Figure 3.**
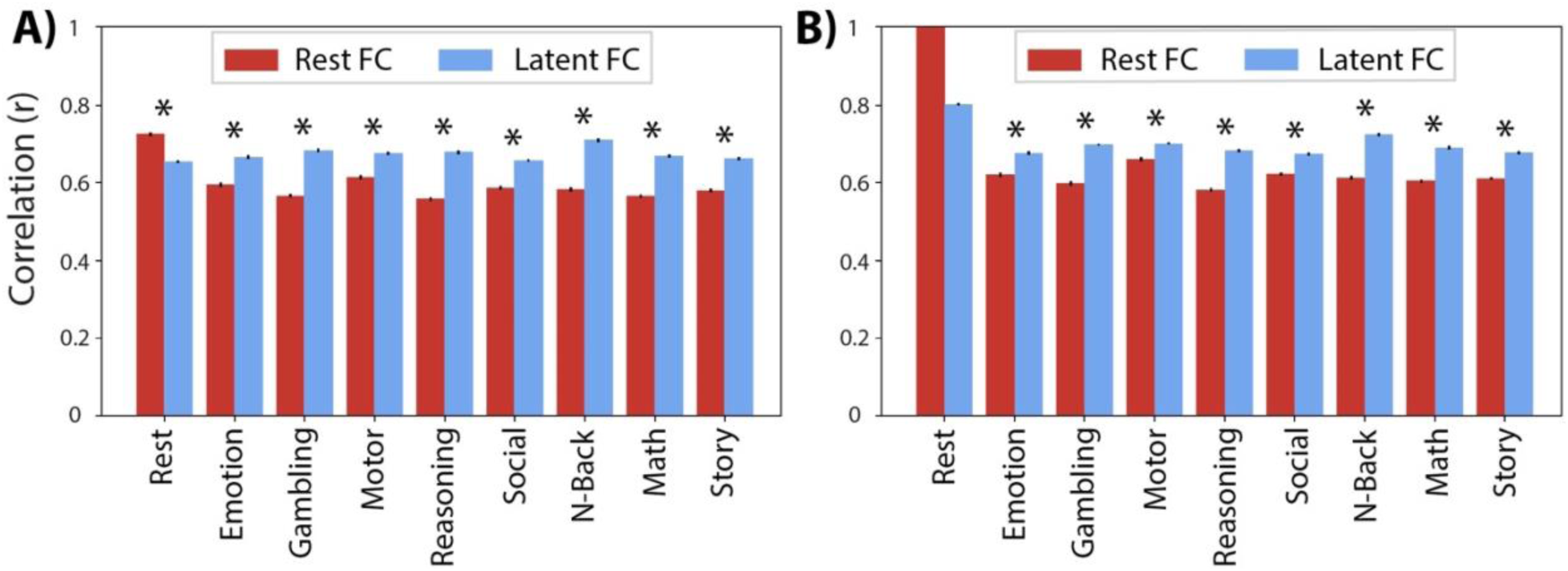
Generalizability of FC patterns. Pearson’s correlation was used to quantify the similarity of latent FC (blue) and rest FC (red) to held-out state FC. Error bars show the standard error of the mean. Asterisks indicate significant differences in similarity of latent FC and rest FC to held-out state FC. **A)** Results when controlling for the number of time points in the resting state data. This included 25 minutes of resting-state fMRI data, matching the total amount of “on-task” time across all tasks. **B)** Results when not controlling for the number of time points (including 58 minutes of resting-state data); the resting state prediction is therefore a perfect reproduction (no error bars or comparison). These results are consistent with resting-state FC overfitting to resting state, reducing its generalizability relative to latent FC.

When using the full timeseries (i.e., not controlling for the amount of data used to obtain the FC estimates across states), we still found greater similarity of latent FC relative to rest FC with the task states. However, latent FC exhibited the greatest similarity to the resting state (*r* = 0.80) and the least similarity to the social task (*r* = 0.67; see Figure 3B). Alongside greater similarity estimates with all states, this suggests that states may converge towards latent FC as we sample substantially more data for any given state (e.g., for resting-state FC, 26 minutes of data per participant were included in the data-restricted analysis vs. 58 minutes of data in the unrestricted analysis). All findings were replicated in the validation dataset (Figure S2).

### Latent FC improves prediction of task activation patterns

We next sought to further test our hypothesis that latent FC is highly generalizable (relative to resting-state FC), this time by testing for generalization beyond FC to patterns of task-evoked activation. We began by using GLMs to estimate the pattern of task-evoked activation for each of 24 task conditions. We then used activity flow mapping (Figure 4A) to predict the pattern of task-evoked activation based on a simple neural network model parameterized using either resting-state FC or latent FC. We used Pearson’s correlation to compute the similarity of predicted-to-actual task activations of two activity flow models with different connectivity estimates based on either latent FC or rest FC. As a global measure of performance, we first correlated the predicted activation patterns from the activity flow model using rest and latent FC with the observed activations. Predicted activation patterns from activity flow models with connectivity based on latent FC (*r* = 0.66) outperformed predictions based on resting state FC (*r* = 0.56) in reproducing the observed beta activation patterns (Figure 4D). We then compared the results of the two models at the region (i.e., prediction for a given region across conditions) and condition (i.e., prediction for a given condition across regions) level.

**Figure 4.**
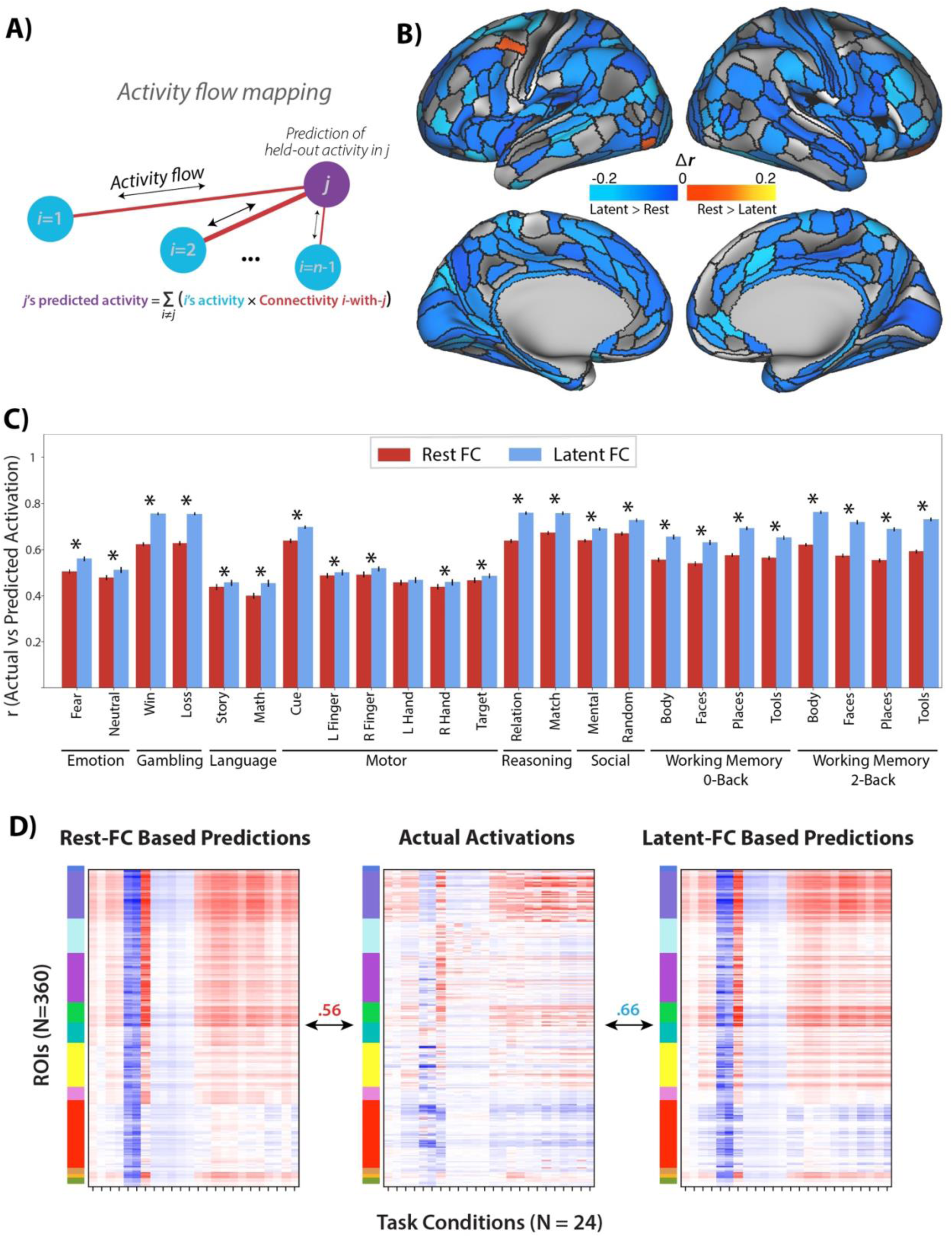
Comparison of activity flow models based on latent FC versus rest FC. **A)** Conceptual model of the activity flow mapping algorithm (Cole et al., 2016) which models the activity of a held-out region (j) as the sum of activity in other brain regions (i) weighted by their shared functional connectivity (ij). **B)** Task activation prediction accuracies by region. Regions with prediction accuracies that were significantly greater using the activity flow model based on latent FC shown in cool colors. Regions with prediction accuracies that were significantly greater using the activity flow model based on rest FC shown in warm colors. The vast majority of regions showing a significant difference showed prediction advantages for latent FC. **C)** Task activation prediction accuracies by condition. Pearson’s correlation was used to quantify the similarity of predicted-to-actual task activations using activity flow models with connectivity based on either rest FC (red) or latent FC (blue). Error bars show the standard error of the mean correlation. Asterisks indicate significant differences in similarity of beta activations from models based on latent FC versus rest FC.

We first estimated predicted beta activations for each region (across conditions) using the activity flow models. This reflects the changes in activation within each region that are dependent on the task condition. For each region, we compared the beta activation predictions of the two activity flow models. For each network, we computed the percent of regions with significantly improved predictions for one of the two models. When using the activity flow model based on latent FC, the predictions were significantly improved (based on a corrected t-test of z-transformed correlation coefficients) for 68% of brain regions (246 out of 360 total), accounting for 33% of VIS1, 69% of VIS2, 64% of SMN, 73% of CON, 70% of DAN, 62% of LAN, 62% of FPN, 100% of AUD, 78% of DMN, 14% of PMM, 0% of VMM, and 33% of ORA. Activity flow based on rest FC significantly improved predictions in 1% of brain regions (4 out of 360 total), accounting for 7% of VIS2 and no other networks (Figure 4B).

When considering prediction accuracy for each task condition, we found that latent FC significantly improved the across-region predicted activations for all task conditions – except the left-hand condition of the motor task – when comparing the relative activations across the topology of the brain within a condition (Figure 4C). Overlap of predicted-to-actual task activations for the activity flow models were variable by task condition. The activity flow model based on latent FC exhibited the greatest similarity to the 2-back body condition of the WM task (*r* = 0.76) and the least similarity to the math condition of the language task (*r* = 0.45). The activity flow model based on rest FC exhibited the greatest similarity to the matching condition of the relational task (*r* = 0.67) and the least similarity to the math condition of the language task (*r* = 0.4). All findings were replicated in the validation dataset (Figure S3).

### Latent FC improves prediction of general intelligence

Our hypothesis that latent FC generalizes better than resting-state FC also predicts that latent FC should be more related to general cognition and behavior, even behavior independent of the particular tasks used for estimating the task-state FC going into the latent FC estimates. We tested whether latent FC improves prediction of general intelligence using psychometric *g* to capture many different behavioral and cognitive measures (Dubois, Galdi, Paul, & Adolphs, 2018; Gottfredson, 1997). We estimated general intelligence (psychometric *g*) using a factor analysis model on behavioral data from a range of cognitive tasks, then tested whether latent FC and/or rest FC measures could predict general intelligence. We combined the exploration and validation samples to increase the number of participants to 352 for this analysis, given the need for additional participants (relative to the other analyses in this study) to achieve reasonable statistical power for individual difference correlations (Yarkoni, 2009). We then employed a multiple linear regression with ridge regularization approach to predict general intelligence from FC. However, one potential confounding issue with simply pooling the full sample data is that the estimated factor scores for latent FC and psychometric-*g* would be influenced by the data of to-be-predicted individuals, introducing circularity into these analyses. To avoid this, we implemented a between-sample cross validation approach. Here, we estimated factor models for latent FC and psychometric-*g* scores in each subsample separately (i.e., exploratory and validation), and predictions for the exploratory subjects were generated from the validation sample regression model and vice versa.

We found that predicted general intelligence was significantly correlated with actual general intelligence for models using both rest FC (*r* = 0.26, *p* = 5.46e-07) and latent FC (*r* = 0.35, *p* = 1.37e-11) (Figure 5A). Consistent with our hypothesis, the model using latent FC significantly improved prediction of general intelligence compared to the model using rest FC (Δ*r* = 0.09, *t* = 1.77, *p* = 0.04, see Eid et al., 2011 for correlation comparison method). The magnitude of this effect was large, as the percent linear variance explained by latent FC (*R^2^* = 0.123) was approximately two times the percent linear variance explained by rest FC (*R^2^* = 0.067). In comparison with the overall sample results, the correlation and difference in *R^2^* was larger for the exploratory sample (Figure 5B) while the validation set showed a more-similar difference in *R^2^* despite lower correlations between predicted and actual psychometric *g* scores for both latent and rest FC data (Figure 5C). A meta-analysis (Field, 2001) of the exploratory and validation samples suggested that the pooled correlation difference effect was significant (Δz_pooled_ = 0.09, *p* = 0.016).

**Figure 5.**
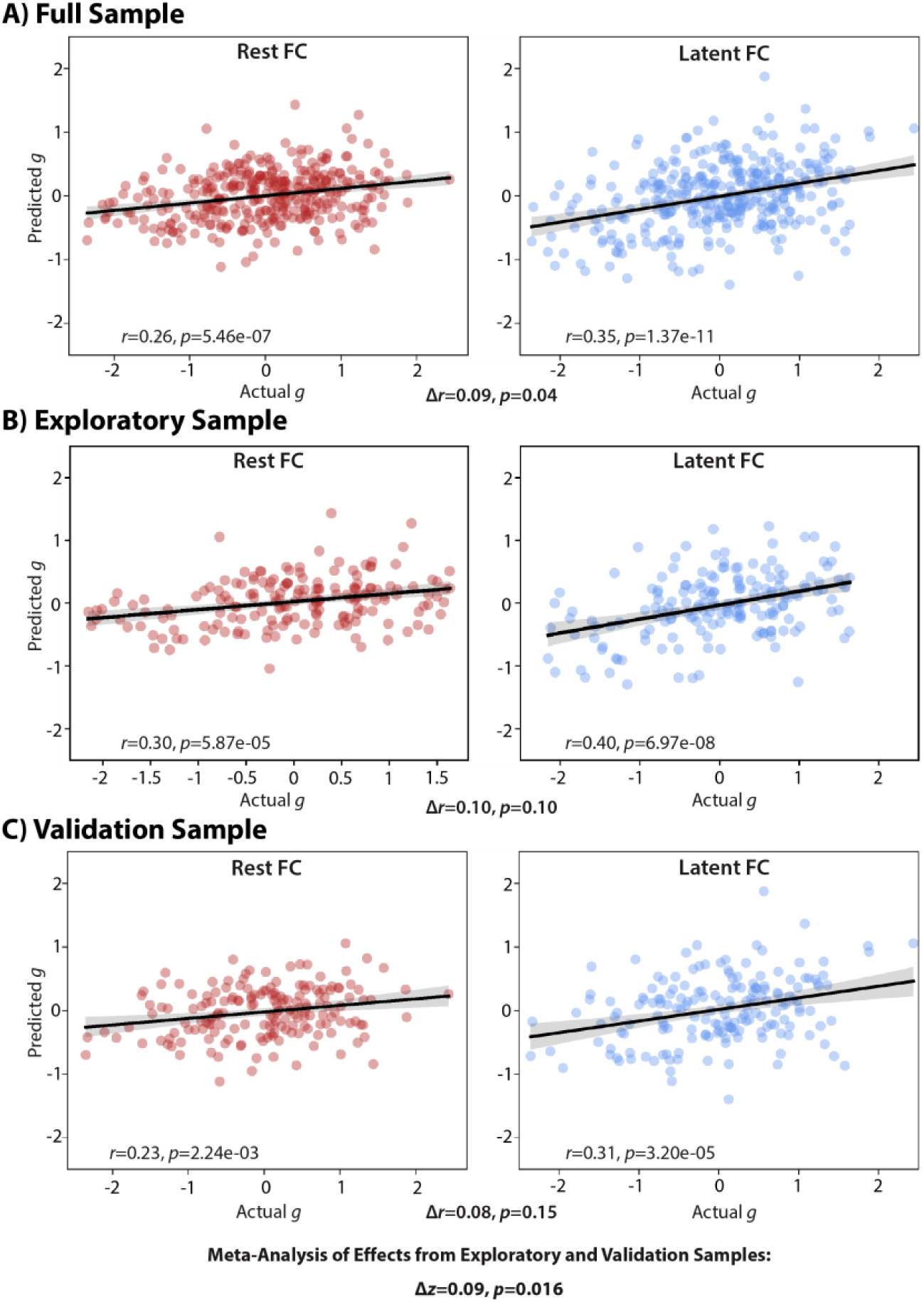
Relationship between actual and predicted general intelligence. Results of ridge regression models from rest FC and latent FC in the **A)** overall, **B)** exploratory, and **C)** validation sample. The significance of the difference in correlation (Δr) is indicated below each plot. A meta-analysis of the exploratory and validation samples showed a significant difference in the correlations between actual and predicted g scores when comparing rest to latent FC (Δz = 0.09, p = 0.016).

### Comparing latent and average FC

While the factor model uses the covariance among the different states to compute optimal weights, a simpler approach to finding consensus among states involves taking a simple average across states. This approach assumes the weights/loadings between measured states are equal. Given that the computed weights in our results with latent FC were relatively uniform across states, we determined that this assumption was reasonable in this case. This supports the use of average FC, however we directly compared latent FC to average FC to assess whether there were any advantages to either method. To compare the factor model with a simple average, we computed the mean value of each edge across states to construct an average connectivity matrix. For all analyses, we controlled for the amount of data between rest and task. Results indicated that combining across states, regardless of the approach, shows substantial improvements over using even the full resting state data. Indeed, the average FC approach appears to out-perform the latent FC approach (albeit only slightly) in generalizing to held-out connectivity states (Figure 6A). In the activity flow mapping results, however, latent FC consistently outperforms average FC in predicting regional activity patterns, showing better predictions in 348 out of 360 regions (97%), whereas average FC showed no improved predictions (Figure 6B). Similarly, latent FC outperformed average FC in condition-wise activity flow predictions in 22 out of 24 conditions (Figure 6C). Together these results suggest that the average FC approach (sometimes termed “general functional connectivity”) is a reasonable alternative to the more complex latent FC approach, so long as the optimal weights across states are close to equal (an assumption not made by latent FC). This difference between the methods would likely become more meaningful in cases wherein a particular brain state is highly distinct from all others (e.g., deep sleep vs. conscious states) or when one or more states is much noisier than the others (which would be weighted lower by latent FC but not average FC).

**Figure 6.**
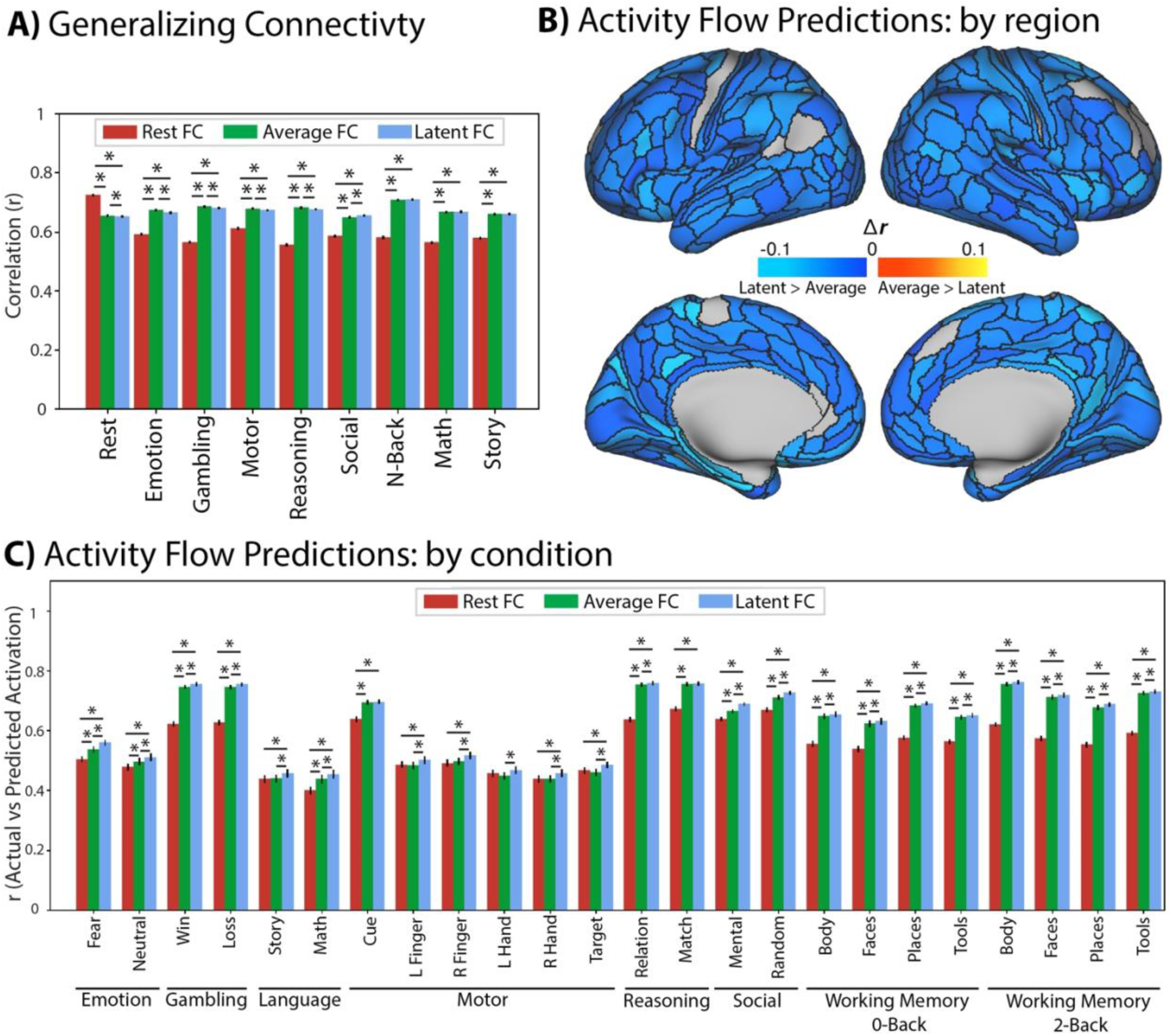
Comparison of A) Generalizing connectivity and B) Activity flow models by region and C) condition, based on latent FC and average FC versus Rest. **A)** Both average (green) and latent (blue) FC outperformed rest FC on generalizing to all task states except resting state. Average FC performed similarly or slightly better than latent FC on all held-out states (asterisks denote significant differences, the higher position represents the test of rest vs latent FC). **B)** In contrast, latent FC outperformed average FC in predicting task activation in 97% of regions, whereas average FC outperformed in 0% regions. **C)** Across all conditions, latent FC was a better predictor than average FC of held-out activations (lower asterisks indicate significant difference between adjacent bars; higher asterisks indicate significant difference between latent and rest FC). All results control for the number of time points in the resting state data. For the validation sample results, see Figure S4.

## Discussion

Defining a map of task-independent, intrinsic functional connections in the brain is a major aim of basic research in cognitive neuroscience. Intrinsic FC persists across task states, making it a more reliable and generalizable measure of the underlying functional dynamics that shape cognition and behavior. As such, measures of intrinsic FC are better candidates to serve as stable biomarkers of important individual differences in behavioral outcomes (Elliott et al., 2019). We utilized a factor analytic approach, a well-developed technique from measurement psychometrics (Bollen, 2002), to define intrinsic FC as a latent variable derived from the common variances in FC across task states. We compared the factor model against the standard approach applied in the field, FC derived from resting state. The factor model not only shows enhanced measurement and predictive properties beyond measures of intrinsic FC derived from resting state, it also offers a unique theoretical perspective on the relationship between intrinsic and task-specific brain states. In a latent variable model, individual task states are viewed as observable sample realizations of the underlying intrinsic connectivity, and task-specific deviations from this baseline are modeled as unique errors arising from a combination of noise and state-specific properties. The factor modeling approach allows researchers to not only gain traction in defining intrinsic FC common among brain states, but also to separate and explore properties that are specific to individuals and states.

### Factor Analytic Model of Functional Connectivity

We began by building factor analysis models of latent FC using two approaches. In the first, we modeled latent FC using all available data. In this model, resting state functional connections had the highest number of significant loadings of any condition. However, when controlling for the number of time-points (by reducing the number of resting state timepoints to match the tasks with shorter durations), resting state connections had the lowest percentage of significant factor loadings. This property of the factor model highlights one of its strengths; higher precision measurements show higher fidelity to the underlying common latent factor than lower-precision measures. Here, the precision appears to be driven primarily by the amount of data. However, in the absence of stringent data quality control, the factor model can also down-weight poor-quality data (e.g., high motion, artifacts) relative to higher-quality data when variability associated with noise does not replicate across task states.

Conversely, tasks that more closely represent underlying intrinsic FC will show stronger factor loadings, similar to how the Raven’s Progressive Matrices task loads highly onto the generalized intelligence factor (Dubois et al., 2018). Given its widespread use as a marker of intrinsic FC, we might have expected that resting state would load highly onto the latent FC factor regardless of how much data went into its estimation. However, when controlling for the amount of data used to estimate FC, the resting state loadings were lower than all other examined states, even though there were still many more TRs of resting state than any one task state. Additionally, when using the full amount of data to estimate rest FC, the factor loadings for resting state was similar to the story and math tasks, each of which were estimated with much less data (Figure 2; values in parentheses). These results suggest that resting state is not an especially good proxy for intrinsic FC, which aligns with its relatively poor performance compared with latent FC in predicting the patterns of connectivity and evoked brain activity observed for other states.

### Latent FC as a Reliable Measure of Intrinsic Connectivity

As mentioned previously, a marker of intrinsic connectivity is its persistence across task states (i.e., generalizability), as well as its ability to accurately recapitulate observed realizations of evoked brain activity and connectivity (Elliott et al., 2019, 2020; Kragel, Han, Kraynak, Gianaros, & Wager, 2020; Parkes, Satterthwaite, & Bassett, 2020). Our results highlight the advantages of latent versus rest FC to reliably predict independent connectivity and regional activations. When comparing patterns of connectivity, we showed that latent FC showed higher correlation with held-out, task-specific connectivity states compared with rest FC, with the sole exception of resting state connectivity where rest FC outperformed. This pattern of results suggests that resting state FC is less generalizable as a measure of intrinsic connectivity and instead there are resting state-specific factors which shape the dynamics of rest FC that are not present in other states.

One potential explanation for this might be that tasks as a group reliably differ from rest FC’s more intrinsic profile, and the reduction in generalizability reflects deviations from a default state. Under this explanation, the latent FC advantage could simply reflect that there are more task indicators in the measurement model than rest (although note that even when controlling for number of timepoints, the amount of rest data is equal to all the tasks combined) and we would predict that latent FC would be a poorer representation of rest FC patterns of connectivity. However, results did not show a substantial drop in the correlation of latent and rest FC compared to the correlations of latent FC with the various task FC patterns (blue bars, Figure 3). Indeed, it is when we used rest FC as the predictor that we observed reductions in its correlation with task connectivity, compared with resting state (red bars, Figure 3). This suggests that latent FC does a better job of representing common, stable variability in FC profiles across both resting and task states. Importantly, latent FC does so even though the task or rest condition being correlated is left out of the factor model for that specific comparison to avoid circularity. As such, the factor score analytically has different indicators across all comparisons, and nevertheless still outperforms rest FC. Moreover, obtaining a better sample of the resting state by using the full time series resulted in the resting state having the highest factor loadings and a strongest correlation with latent FC, which suggests that over time the resting state converges to latent FC.

The advantages of latent FC are not, however, restricted to the connectivity space; the latent measure of intrinsic FC also outperforms rest FC in predicting state-specific activation patterns. Not only did latent FC support higher prediction accuracy by the activity flow model of task activation globally (*r_latent_* = 0.66 vs. *r_rest_* = 0.56), it showed condition-specific advantages in 23 out of 24 specific task conditions (Figure 4B). Rest FC, in comparison, displayed higher prediction accuracy in none of the task conditions (in the left-hand motor condition, latent and rest FC performed comparably; Figure 4C). When we examined predictions of region-specific patterns of activation, results showed that latent FC had improved prediction over rest FC for 68% of all brain regions across a variety of distributed networks. In contrast, rest FC showed improved prediction for only 1.1% of regions, all of which were restricted to the VIS2 network (and constituted only 7% of that network). These improvements, as before, were not due to circularity in the analyses, as task predictions using latent FC were done using the leave-one-task-out approach in the factor model.

### Improving External Validity with Latent FC

While latent FC has demonstrable advantages for prediction within the brain, its utility as a method of estimating brain-based biomarkers relies on its predictive validity for outcomes of interest. Here, we showed that connectivity values from latent FC showed superior prediction of a metric of generalized intelligence (psychometric *g*) than did rest FC connections. Although both rest FC and latent FC values significantly predicted individual differences in generalized intelligence, latent FC nearly doubled the percent of explained variance in the outcome over rest FC (∼12% versus ∼7%). In measurement science, this is a hallmark advantage of the latent variable approach used in factor analysis. Methods which fail to account for measurement error tend to show reduced relationships between variables, whereas modeling state-specific error terms disattenuates those relationships (Schmidt & Hunter, 1996). Indeed, generalized intelligence is generally modeled with a factor analytic approach for precisely this reason. We demonstrate that the framework for improving measurement properties in behavioral measures applies equally to measures derived from functional neuroimaging data. As such, factor analytic models are ideal for aiding the search for biomarkers across a wide domain of individual difference outcomes. Furthermore, more reliable estimates of FC may aid modeling efforts that use intermediate network metrics (e.g., modularity, hub diversity) to predict participant behavior (e.g., (Bertolero, Yeo, Bassett, & D’Esposito, 2018)), offer an exciting range of possible uses for latent FC in future work.

### State Aggregation Improves Predictive Performance

The performance of average FC suggests that aggregating information across states has advantages over longer scan sessions of resting state, regardless of the approach used. Interestingly, average FC performance is not uniform in relation to the latent FC, performing as-good or slightly better than latent FC in correlating with state-specific connectivity, but underperforming latent FC in predicting held out activity in almost all regions. A few circumstances may predict when we would expect to see more or less pronounced differences between average and latent FC. First, data quality: We expect more pronounced differences for lower quality data and less pronounced differences for higher quality data. The HCP data used here is of extremely high quality, which reduces variability in noise between scans. This is reflected in the average factor loadings which are relatively close in value across states (Figure 2). Of course, as the loadings converge in value, the more similar average and latent FC will become (here the connectivity values are correlated; *r* = .98). Second, the method of factor analysis used: Here, we opted to fit a single-factor model for each connection independently due to the large number of operations (e.g., separate models for each connection). However, a single factor in isolation may not be the best fit for brain data (van Kesteren & Kievit, 2020) and the method here might represent a sort of floor performance for latent FC relative to approaches which adopt a dependent model that tries to optimize the fit for each factor model.

Finally, there appear to be differences depending on the type of dependent variable in question. For example, while the factor and average models converge in their correlation with connectivity for held-out states, we found that activity flow models that incorporated latent FC performed better. Average and factor models produced similar patterns of relative connectivity (i.e., highly correlated patterns of FC), however the distribution of connectivity values differ. Latent FC estimates exhibited a sparser distribution of connectivity by zero-ing out low and/or unstable connections, which may have improved the activity flow models by reducing the contributions of disconnected brain regions (see Figure S5).

Despite the relatively small differences in performance between average and latent FC, there are theoretical reasons to prefer a latent variable perspective for FC estimation. The first, as mentioned before, is that while the average FC must assume equal loadings, latent FC makes this a testable hypothesis. If loadings converge towards equal values, then average and latent FC will converge (as they nearly did here). This suggests that averaging will likely perform well under conditions similar to the HCP data (high quality, young adult data). However, as the data diverges from this baseline, latent FC should have advantages by weighting data according to how closely it reflects intrinsic functional states and contributes to the common variance across measures. If differences among measures increase (i.e., measures reflect intrinsic FC better or worse), we would hypothesize that average and latent FC would diverge in their performance. We can see this in a small reproducible example (see Supplemental Code Demonstrations), where more variable loadings impact the ability of sums scores, but not factor scores, to predict a hypothetical outcome variable. However, apart from these practical considerations, a latent variable model of FC is a good theoretical model for how state-specific functional connections emerge from underlying, intrinsic neural connectivity. Intrinsic connectivity is an unobserved state (Bollen, 2002) that gives rise to state specific phenotypes based on combinations of common (i.e., the latent factor) and state-specific (i.e., the error) variance.

## Conclusions

In summary, we utilized a factor analytic approach to derive intrinsic FC from multiple task and resting state data. Our derived measure, termed latent FC, showed improved generalizability and reliability compared to a standard measure of resting-state FC. Not only did latent FC do a better job of reflecting state-specific FC patterns across tasks, it also overwhelmingly improved predictions of regional activations when utilized in activity flow models. Finally, connectivity derived from latent FC doubled the predictive utility of an external measure of generalized intelligence (*g*) compared with connectivity from rest FC, highlighting its suitability for use in clinical and other individual difference research, where reliable biomarkers are needed. These results present compelling support for the use of factor analytic models in cognitive neuroscience, demonstrating the value of established tools from psychometrics for enhancing measurement quality in neuroscience.

## Materials and Methods

For clarity, portions of the text in this section are from our prior publication using the same dataset and some identical analysis procedures: Ito et al. (2020).

### Participants

Data in the present study were collected as part of the Washington University-Minnesota Consortium of the Human Connectome Project (HCP) (Van Essen et al., 2013). A subset of data (n = 352) from the HCP 1200 release was used for empirical analyses. Specific details and procedures of subject recruitment can be found in Van Essen et al. (2020). The subset of 352 participants was selected based on: quality control assessments (i.e., any participants with any quality control flags were excluded, including 1) focal anatomical anomaly found in T1w and/or T2w scans, 2) focal segmentation or surface errors, as output from the HCP structural pipeline, 3) data collected during periods of known problems with the head coil, 4) data in which some of the FIX-ICA components were manually reclassified; exclusion of high-motion participants (participants that had any fMRI run in which more than 50% of TRs had greater than 0.25mm framewise displacement); removal according to family relations (unrelated participants were selected only, and those with no genotype testing were excluded). A full list of the 352 participants used in this study will be included as part of the code release.

All participants were recruited from Washington University in St. Louis and the surrounding area. We split the 352 subjects into two cohorts of 176 subjects: an exploratory cohort (99 women) and a validation cohort (84 women). The exploratory cohort had a mean age of 29 years of age (range=22-36 years of age), and the validation cohort had a mean age of 28 years of age (range=22-36 years of age). All subjects gave signed, informed consent in accordance with the protocol approved by the Washington University institutional review board.

### Scan Acquisition

Whole-brain multiband echo-planar imaging acquisitions were collected on a 32-channel head coil on a modified 3T Siemens Skyra with TR=720 ms, TE=33.1 ms, flip angle=52°, Bandwidth=2,290 Hz/Px, in-plane FOV=208180 mm, 72 slices, 2.0 mm isotropic voxels, with a multiband acceleration factor of 8. Data for each subject were collected over the span of two days. On the first day, anatomical scans were collected (including T1-weighted and T2-weighted images acquired at 0.7 mm isotropic voxels) followed by two resting-state fMRI scans (each lasting 14.4 minutes) and ending with a task fMRI component. The second day consisted of first collecting a diffusion imaging scan, followed by a second set of two resting-state fMRI scans (each lasting 14.4 minutes), and again ending with a task fMRI session.

Each of the seven tasks was collected over two consecutive fMRI runs. The seven tasks consisted of an emotion cognition task, a gambling reward task, a language task, a motor task, a relational reasoning task, a social cognition task, and a working memory task. Briefly, the emotion cognition task required making valence judgements on negative (fearful and angry) and neutral faces. The gambling reward task consisted of a card guessing game, where subjects were asked to guess the number on the card to win or lose money. The language processing task consisted of interleaving two language conditions, which involved answering questions related to a story presented aurally, and a math condition, which involved basic arithmetic questions presented aurally. Note that we treated the two language task conditions as separate tasks, given the highly distinct nature of the conditions (other than that they were presented aurally). The motor task involved asking subjects to either tap their left/right fingers, squeeze their left/right toes, or move their tongue. The reasoning task involved asking subjects to determine whether two sets of objects differed from each other in the same dimension (e.g., shape or texture). The social cognition task was a theory of mind task, where objects (squares, circles, triangles) interacted with each other in a video clip, and subjects were subsequently asked whether the objects interacted in a social manner. Lastly, the working memory task was a variant of the N-back task. Further details on the resting-state fMRI portion can be found in (Smith et al., 2013), and additional details on the task fMRI components can be found in (Barch et al., 2013).

### Behavior: Data

To assess generalized intelligence (*g*), we drew 11 measures of cognitive ability from the HCP dataset, which are derived from the NIH Toolbox for Assessment of Neurological and Behavioral function (http://www.nihtoolbox.org; (Gershon et al., 2013) and the Penn computerized neurocognitive battery (Gur et al., 2010). Tasks included: picture sequence memory; dimensional card sort; flanker attention and inhibitory control; the Penn Progressive Matrices; oral reading recognition; picture vocabulary; pattern completion processing speed; variable short Penn line orientation test; Penn word memory test (number correct and median reaction time as separate variables]) and list sorting. For all measures, the age-unadjusted score was used where applicable. For complete information regarding all measures, see the descriptions in the Cognition Category of the HCP Data Dictionary (https://wiki.humanconnectome.org/display/PublicData/HCP+Data+Dictionary+Public-+Updated+for+the+1200+Subject+Release).

### Behavior: Factor analysis model of psychometric ‘g’

We then derived a general factor of intelligences using a multiple-indicator latent factor model. We approach the factor model using a confirmatory factor analysis (CFA) approach with a unitary factor underlying all individual cognitive tasks. Factor loadings were estimated using the *psych* R package (Revelle, 2017). Factor scores were computed using the regression method (Thurstone, 1935) to obtain manifest variables for prediction.

### fMRI: Preprocessing

Minimally preprocessed data for both resting-state and task fMRI were obtained from the publicly available HCP data. Minimally preprocessed surface data was then parcellated into 360 brain regions using the Glasser atlas (Glasser et al., 2016). We performed additional preprocessing steps on the parcellated data for resting-state fMRI and task state fMRI to conduct neural variability and FC analyses. This included removing the first five frames of each run, de-meaning and de-trending the time series, and performing nuisance regression on the minimally preprocessed data (Ciric et al., 2017). Nuisance regression removed motion parameters and physiological noise. Specifically, six primary motion parameters were removed, along with their derivatives, and the quadratics of all regressors (24 motion regressors in total). Physiological noise was modeled using aCompCor on time series extracted from the white matter and ventricles (Behzadi, Restom, Liau, & Liu, 2007). For aCompCor, the first 5 principal components from the white matter and ventricles were extracted separately and included in the nuisance regression. In addition, we included the derivatives of each of those components, and the quadratics of all physiological noise regressors (40 physiological noise regressors in total). The nuisance regression model contained a total of 64 nuisance parameters. This was a variant of previously benchmarked nuisance regression models reported in (Ciric et al., 2017).

We excluded global signal regression (GSR), given that GSR can artificially induce negative correlations (Murphy, Birn, Handwerker, Jones, & Bandettini, 2009; Power et al., 2014), which could bias analyses of whether global correlations decrease during task performance. We included aCompCor as a preprocessing step here given that aCompCor does not include the circularity of GSR (regressing out some global gray matter signal of interest) while including some of the benefits of GSR (some extracted components are highly similar to the global signal) (Power et al., 2018). This logic is similar to a recently-developed temporal-ICA-based artifact removal procedure that seeks to remove global artifact without removing global neural signals, which contains behaviorally relevant information such as vigilance (Glasser et al., 2018; Wong, Olafsson, Tal, & Liu, 2013). We extended aCompCor to include the derivatives and quadratics of each of the component time series to further reduce artifacts. Code to perform this regression is publicly available online using python code (version 2.7.15) (https://github.com/ito-takuya/fmriNuisanceRegression). Following nuisance regression, the time series for each run (task-state and rest-state) were z-normalized such that variances across runs would be on the same scale (i.e., unit variance).

Task data for task FC analyses were additionally preprocessed using a standard general linear model (GLM) for fMRI analysis. For each task paradigm, we removed the mean evoked task-related activity for each task condition by fitting the task timing (block design) for each condition using a finite impulse response (FIR) model (Cole et al., 2019). (There were 24 task conditions across seven cognitive tasks.) We used an FIR model instead of a canonical hemodynamic response function given recent evidence suggesting that the FIR model reduces both false positives and false negatives in the identification of FC estimates (Cole et al., 2019). This is due to the FIR model’s ability to flexibly fit the mean-evoked response across all blocks.

FIR modeled task blocks were modeled separately for task conditions within each of the seven tasks. In particular, two conditions were fit for the emotion cognition task, where coefficients were fit to either the face condition or shape condition. For the gambling reward task, one condition was fit to trials with the punishment condition, and the other condition was fit to trials with the reward condition. For the language task, one condition was fit for the story condition, and the other condition was fit to the math condition. For the motor task, six conditions were fit: (1) cue; (2) right hand trials; (3) left hand trials; (4) right foot trials; (5) left foot trials; (6) tongue trials. For the relational reasoning task, one condition was fit to trials when the sets of objects were matched, and the other condition was fit to trials when the objects were not matched. For the social cognition task, one condition was fit if the objects were interacting socially (theory of mind), and the other condition was fit to trials where objects were moving randomly. Lastly, for the working memory task, 8 conditions were fit: (1) 2-back body trials; (2) 2-back face trials; (3) 2-back tool trials; (4) 2-back place trials; (5) 0-back body trials; (6) 0-back face trials; (7) 0-back tool trials; (8) 0-back place trials. Since all tasks were block designs, each time point for each block was modeled separately for each task condition (i.e., FIR model), with a lag extending up to 25 TRs after task block offset.

### fMRI: Task activation

We performed a standard task GLM analysis on fMRI task data to estimate evoked brain activity during task states. The task timing for each of the 24 task conditions was convolved with the SPM canonical hemodynamic response function to obtain task-evoked activity estimates (Friston et al., 1994). Coefficients were obtained for each parcel in the Glasser et al. (2016) cortical atlas for each of the 24 task conditions.

### fMRI: Functional connectivity (FC) estimation

Residual timeseries from the rest and task nuisance regressions were used to estimate functional connectivity for each task. Connectivity values were estimated using zero-lag Pearson product-moment correlations. Timeseries were concatenated across separate runs of the same task to yield a single connectivity value per edge for a given task or resting state condition. For each task scan, we utilized TRs that corresponded to “on-task” timepoints. For instance, we extracted TRs from the working memory scan during N-back task blocks, excluding TRs from the inter-block fixation periods. For the number of TRs included in the connectivity estimates for each condition and scan state, see Table S1.

### fMRI: Factor analysis model of latent FC

Factor analysis for obtaining latent FC was conducted with the same approach used to obtain factor scores for generalized intelligence. FC estimates from each separate fMRI task were used as indicators on a unitary factor model and factor scores were obtained using the regression method in the *psych* R package. A separate model was computed for each edge in the connectivity adjacency matrix. We took several approaches to test the predictive utility of latent FC for activation and behavior (detailed below).

The first set of analyses tested two alternative measurement approaches for latent FC. The first was to utilize all available data from each functional scan to estimate factor scores for each edge. However, because of the differential amount of scan time for different functional runs (e.g., ∼58 minutes of resting-state versus ∼10 minutes of working memory scans), we might expect indicators (i.e., scan types) with more data would dominate the measurement model in the factor analysis. To control for this potential confound, we ran additional analyses where indicators were constrained to have equivalent numbers of TRs used to estimate individual scan functional edges between task and rest, and between different task states. The reasoning task had the fewest “on-task” TRs (264) and therefore served as the limiting factor for task scans data. As such, 264 TRs of each task (for 2112 TRs of task) and a corresponding 2112 TRs of rest were used in these analyses. All of these analyses were performed modeling all available scan types in the same factor model.

For activity flow mapping (ActFlow) analyses (Cole et al., 2016; Ito et al., 2020), where activations in held-out regions were predicted using estimated activity flowing over estimated connections, latent FC was estimated independently for each connection by applying leave-one-state-out factor analysis (LOSO-FA) on the state FC estimates to prevent circularity in the predictive model. For instance, when predicting activation in the emotion task, FC estimates were obtained without including the emotion task as an indicator in the factor model. In all ActFlow analyses, we estimated predictions per subject and then pooled results (i.e., an estimate-then-average approach).

### Meta-Analysis Across Samples

To appropriately combine effects of the *g*-prediction analysis across the validation and exploratory sample, we computed r-to-z-score transformations of the individual coefficients and then combined them into a weighted z score using the standard formula (Field, 2001) where *z* is the z-score and the weight (*w*) corresponds to the sample size.

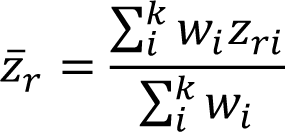

The significance of this meta-analytic parameter was determined using a chi-square test (Field, 2001).

## Acknowledgements

The authors acknowledge support by the US National Institutes of Health under awards R01 AG055556 and R01 MH109520 to MWC. Data were provided by the Human Connectome Project, WU-Minn Consortium (Principal Investigators: D. Van Essen and K. Ugurbil; 1U54MH091657) funded by the 16 NIH Institutes and Centers that support the NIH Blueprint for Neuroscience Research; and by the McDonnell Center for Systems Neuroscience at Washington University. The authors acknowledge the Office of Advanced Research Computing (OARC) at Rutgers, The State University of New Jersey for providing access to the Amarel cluster and associated research computing resources that have contributed to the results reported here. The content is solely the responsibility of the authors and does not necessarily represent the official views of any of the funding agencies.

## Author contributions

KLA and MWC designed the analytic approach. EMM and KLA carried out analyses with MWC. EMM, KLA, and MWC wrote the manuscript, with feedback received from all other authors.

