## Supplemental Material for "Latent functional connectivity underlying multiple brain states"

| | All Data | Controlling for # Timepoints |
|---|---|---|
| **State** | **# TR** | |
| Rest | 4800 | 2112 |
| Emotion | 274 | 264 |
| Gambling | 304 | 264 |
| Motor | 384 | 264 |
| Reasoning | 264 | 264 |
| Social | 310 | 264 |
| Working Memory | 608 | 264 |
| Math | 311 | 264 |
| Language | 284 | 264 |

***Table S1. Number of TRs used in FC estimation per task and condition.*** *We utilized all "on-task" TRs (i.e., excluding fixation periods) in the All Data condition. However, to control for differences between task states we equated the number of TRs (264) for each. To balance task and rest information in the full model, we then drew TRs from the resting state data to be equal to the total number of task TRs (2112).*

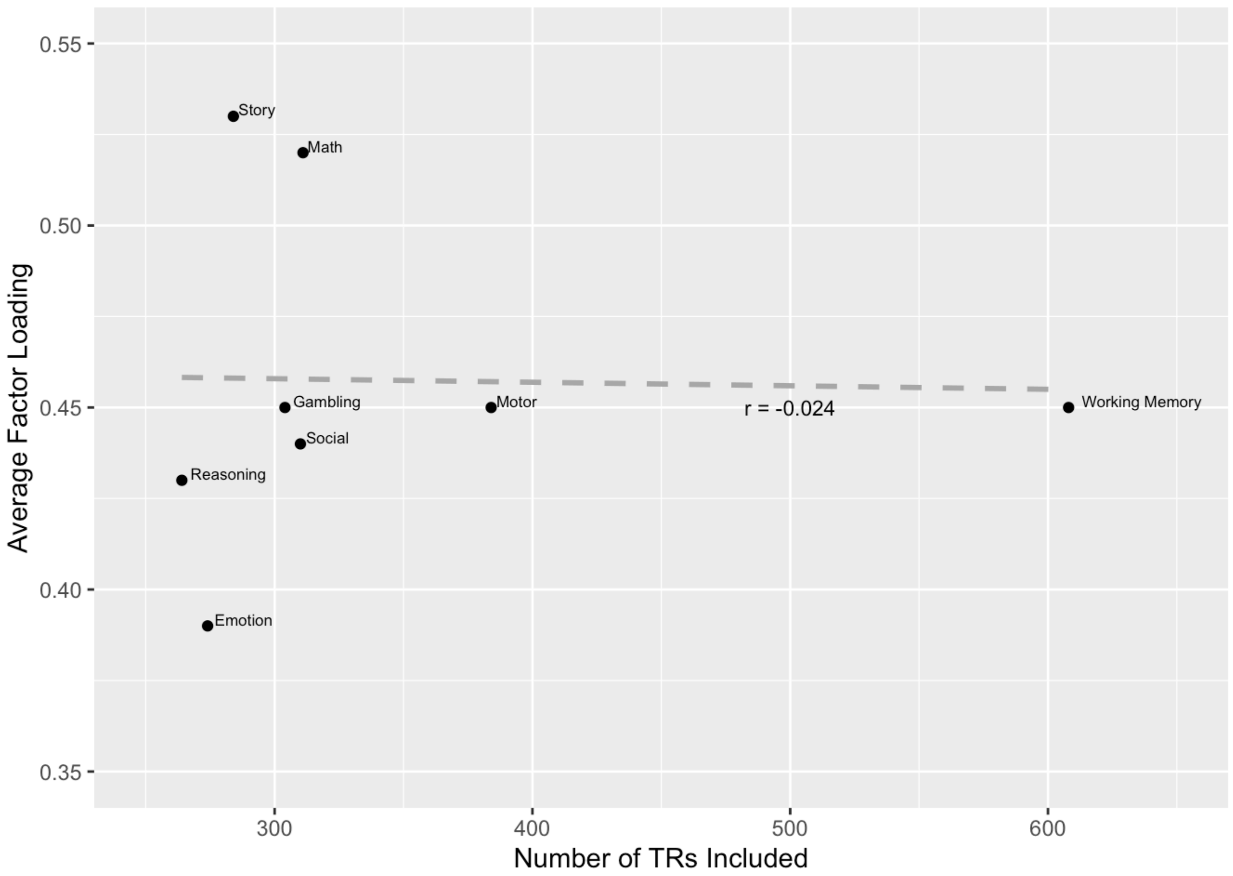


***Figure S1: Number of TRs included in the Latent FC estimate and the Average Factor Loading.*** *Even though different task FC states were characterized different numbers of TRs, there was no relationship between this number of available TRs and the average factor loading in the model. This suggests that the strength of the relationship of each task indicator to the Latent FC estimate was not driven simply by the amount of data available.*

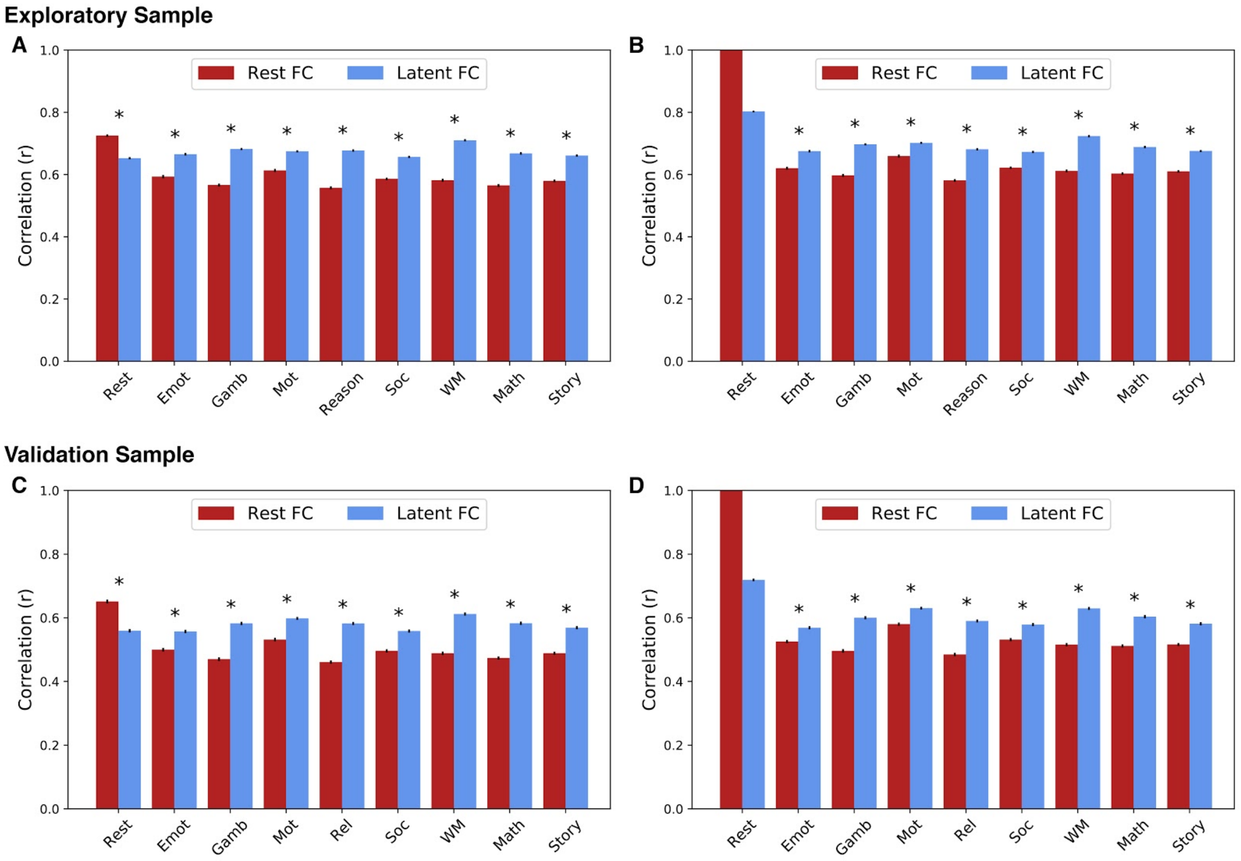


***Figure S2: Generalizability of FC patterns for Exploratory and Validation Samples.*** *Pearson's correlation was used to quantify the similarity of latent FC (blue) and rest FC (red) to held-out state FC. Error bars show the standard error of the mean. Asterisks indicate significant differences in similarity of latent FC and rest FC to held-out state FC.* ***A & C.*** *Results when controlling for the number of time points in the resting state data.* ***B & D.*** *Results when* ***not*** *controlling for the number of time points; the resting state prediction is therefore a perfect reproduction (no error bars or comparison). The amount of resting-state data per participant went from 4800 TRs (58 minutes) to 2112 TRs (25 minutes) when matching the total amount of "on-task" time. These results are consistent with resting-state FC overfitting to resting state, reducing its generalizability relative to latent FC. The overall pattern of results was consistent across samples.*

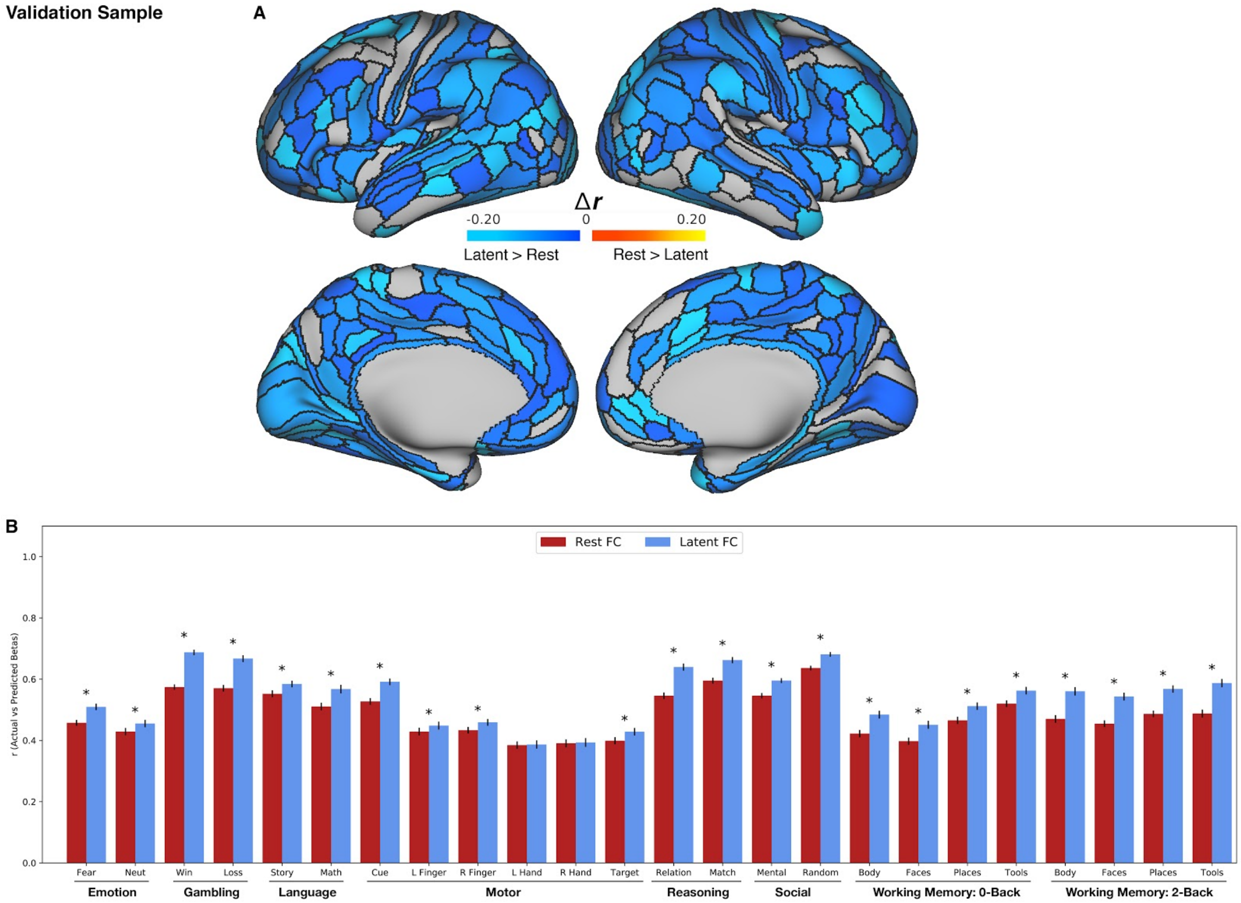


***Figure S3: Comparison of activity flow models based on latent FC versus rest FC: Validation Sample. A.*** *Task activation prediction accuracies by region. Regions with prediction accuracies that were significantly greater using the activity flow model based on latent FC shown in cool colors. Regions with prediction accuracies that were significantly greater using the activity flow model based on rest FC shown in warm colors. The vast majority of regions showing a significant difference showed prediction advantages for latent FC, and no regions showed prediction advantage for rest FC.* ***B.*** *Task activation prediction accuracies by condition. Pearson's correlation was used to quantify the similarity of predicted-to-actual beta activations using activity flow models with connectivity based on either rest FC (red) or latent FC (blue). Error bars show the standard error of the mean correlation. Asterisks indicate significant differences in similarity of beta activations from models based on latent FC versus rest FC.*

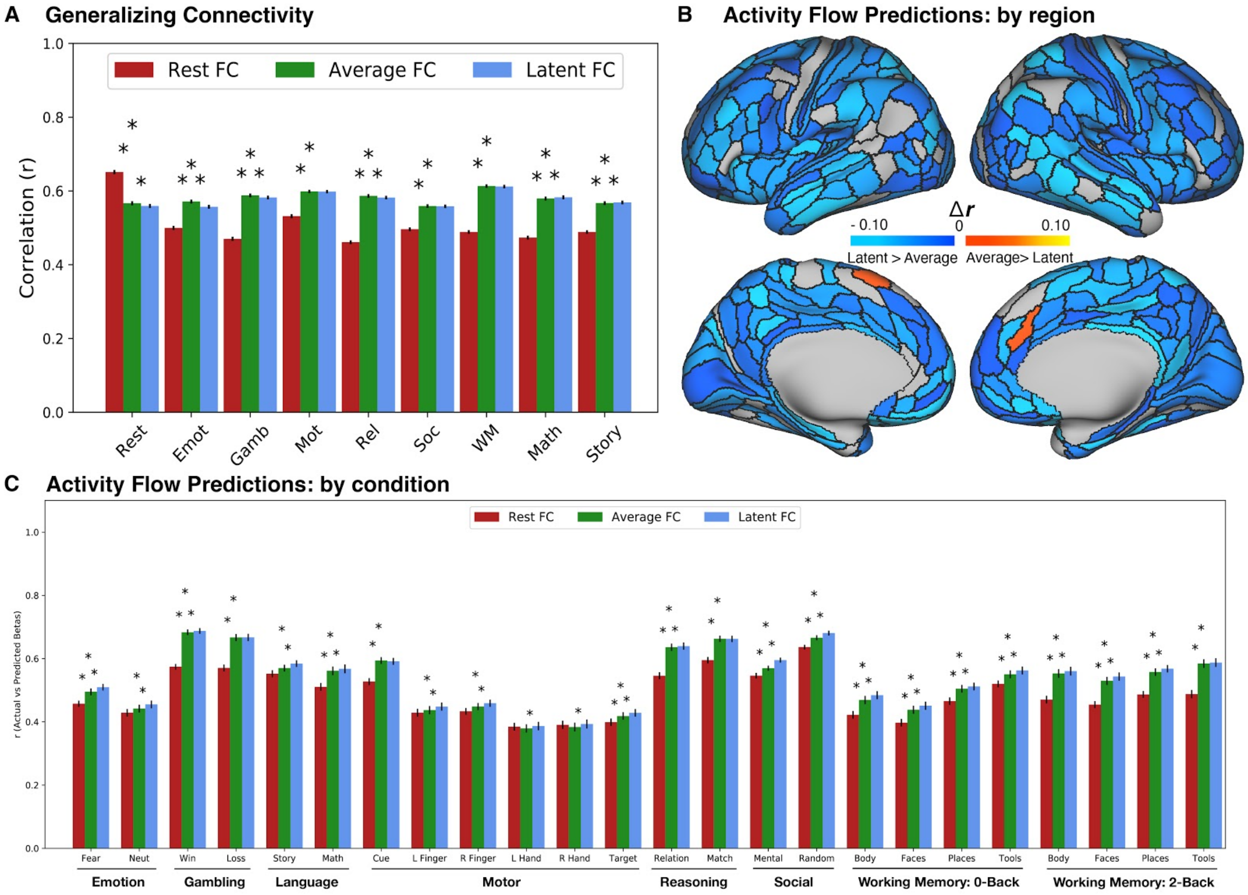


***Figure S4: Comparison of A) Generalizing connectivity and B) Activity flow models by region and C) condition, based on latent FC and average FC versus Rest: Validation Sample. A.*** *Both average (green) and latent (blue) FC outperformed rest FC on generalizing to all task states except resting state. Average FC performed similarly or slightly better than latent FC on all held-out states (asterisks denote significant differences, the higher position represents the test of rest vs latent FC).* ***B.*** *In contrast, latent FC outperformed average FC in predicting task activation in 95% of regions, whereas average FC outperformed in 0.6% regions.* ***C.*** *Across all conditions, latent FC was a better predictor than average FC of held-out activations (lower asterisks indicate significant difference between adjacent bars; higher asterisks indicate significant difference between latent and rest FC).*

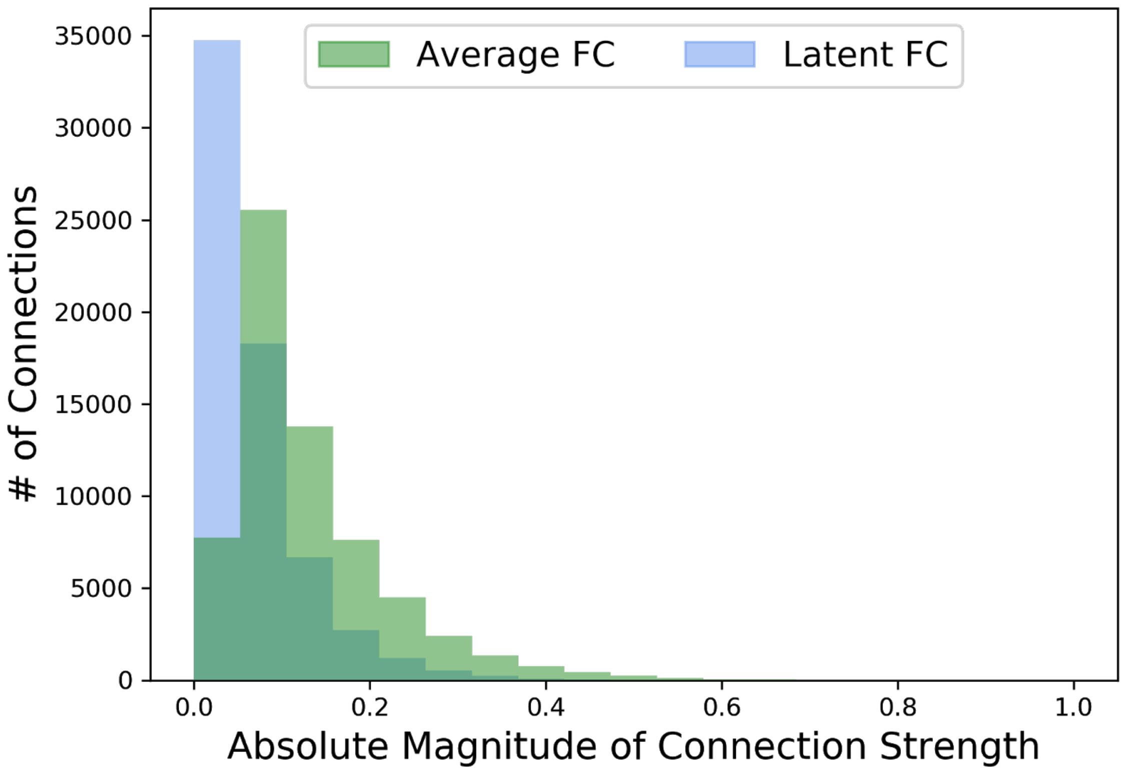


***Figure S5: Distribution of Latent and Average FC Scores.*** *While highly correlated (r = 0.98), latent FC (blue) produces greater sparsity in the connectivity matrix (i.e., more values at or near 0) compared with average FC (green).*

### *Reproducible Example: Impact of Variable Standard Factor Loadings on Average and Factor Model*

#### *Low Standardized Loading Variance*

```
set.seed(1234567)
Nsubs = 500
loadings = runif(9, 0.45, 0.55)

dat01 = data.frame(eta = rnorm(Nsubs, 0, 1)) %>%
  mutate(out = eta) %>%
  mutate(out = out + rnorm(Nsubs, mean=0, sd=1))
for (l in (1:length(loadings))){
  dat01 = dat01 %>%
    mutate(x = 0 + loadings[l]*eta) %>%
    mutate(x = x + rnorm(Nsubs, mean=0, sd=sd(x))) %>%
    plyr::rename(c('x' = paste0('x',l)))
}
dat01 = dat01 %>%
  mutate(sum = rowMeans(dat01[,3:11]))

mod01 = 'factor =~ NA*x1 + x1 + x2 + x3 + x4 + x5 + x6 + x7 + x8 + x9
         factor ~ 0*1
         factor ~~ 1*factor'
fit01 = sem(mod01, data=dat01, estimator='ML')

## lavaan 0.6-9 ended normally after 17 iterations
##
##   Estimator                                         ML
##   Optimization method                           NLMINB
##   Number of model parameters                        27
##
##   Number of observations                           500
##
## Model Test User Model:
##
##   Test statistic                                35.665
##   Degrees of freedom                                27
##   P-value (Chi-square)                           0.123
##
## Parameter Estimates:
##
##   Standard errors                             Standard
##   Information                                 Expected
##   Information saturated (h1) model          Structured
##
## Latent Variables:
##                    Estimate  Std.Err  z-value  P(>|z|)   Std.lv  Std.all
##   factor =~
##     x1                0.498    0.030   16.689    0.000    0.498    0.682
##     x2                0.498    0.028   17.860    0.000    0.498    0.718
##     x3                0.528    0.029   17.955    0.000    0.528    0.720
##     x4                0.466    0.026   17.810    0.000    0.466    0.716
##     x5                0.502    0.029   17.570    0.000    0.502    0.709
##     x6                0.501    0.027   18.248    0.000    0.501    0.729
##     x7                0.415    0.024   17.423    0.000    0.415    0.705
```

```
##     x8                 0.511    0.029   17.730    0.000    0.511    0.714
##     x9                 0.469    0.028   16.876    0.000    0.469    0.688
##
## Intercepts:
##                    Estimate  Std.Err  z-value  P(>|z|)   Std.lv  Std.all
##     factor             0.000                               0.000    0.000
##    .x1                -0.018    0.033   -0.554    0.580   -0.018   -0.025
##    .x2                 0.037    0.031    1.200    0.230    0.037    0.054
##    .x3                 0.048    0.033    1.469    0.142    0.048    0.066
##    .x4                 0.031    0.029    1.055    0.291    0.031    0.047
##    .x5                -0.002    0.032   -0.053    0.958   -0.002   -0.002
##    .x6                 0.019    0.031    0.610    0.542    0.019    0.027
##    .x7                 0.023    0.026    0.861    0.389    0.023    0.039
##    .x8                -0.002    0.032   -0.071    0.943   -0.002   -0.003
##    .x9                 0.036    0.031    1.183    0.237    0.036    0.053
##
## Variances:
##                    Estimate  Std.Err  z-value  P(>|z|)   Std.lv  Std.all
##     factor             1.000                               1.000    1.000
##    .x1                 0.285    0.020   14.294    0.000    0.285    0.534
##    .x2                 0.234    0.017   13.960    0.000    0.234    0.485
##    .x3                 0.258    0.019   13.930    0.000    0.258    0.481
##    .x4                 0.207    0.015   13.975    0.000    0.207    0.487
##    .x5                 0.249    0.018   14.049    0.000    0.249    0.497
##    .x6                 0.221    0.016   13.834    0.000    0.221    0.469
##    .x7                 0.174    0.012   14.092    0.000    0.174    0.503
##    .x8                 0.252    0.018   14.000    0.000    0.252    0.491
##    .x9                 0.245    0.017   14.245    0.000    0.245    0.527
dat01 = dat01 %>%
  mutate(factor.score = lavPredict(fit01))

low.var.sum01 = lm(scale(out) ~ scale(sum), data=dat01)
low.var.factor01 = lm(scale(out) ~ scale(factor.score), data=dat01)
```

**Low Standardized Factor Loading Variance**

| | **Outcome ~ Sum Score** | | | **Outcome ~ Factor Score** | | |
|---|---|---|---|---|---|---|
| *Predictors* | *Estimates* | *SE* | *P-Value* | *Estimates* | *SE* | *P-Value* |
| Intercept | -0.000 | 0.034 | 1.000 | -0.000 | 0.034 | 1.000 |
| Sum Score | 0.653 | 0.034 | **<0.001** | | | |
| Factor Score | | | | 0.652 | 0.034 | **<0.001** |
| Observations | 500 | | | 500 | | |
| $R^2$ / $R^2$ adjusted | 0.427 / 0.426 | | | 0.426 / 0.425 | | |

***High Standardized Loading Variance***

```
set.seed(1234567)
Nsubs = 500
loadings = runif(9, 0.45, 0.55)

dat02 = data.frame(eta = rnorm(Nsubs, 0, 1)) %>%
  mutate(out = eta) %>%
  mutate(out = out + rnorm(Nsubs, mean=0, sd=sd(out)))
for (l in (1:length(loadings))){
  dat02 = dat02 %>%
    mutate(x = 0 + loadings[l]*eta) %>%
    mutate(x = x + rnorm(Nsubs, mean=0, sd=runif(1, .5,3)*sd(x))) %>%
    plyr::rename(c('x' = paste0('x',l)))
}
dat02 = dat02 %>%
  mutate(sum = rowMeans(dat02[,3:11]))

mod02 = 'factor =~ NA*x1 + x1 + x2 + x3 + x4 + x5 + x6 + x7 + x8 + x9
         factor ~ 0*1
         factor ~~ 1*factor'
fit02 = sem(mod02, data=dat02, estimator='ML')

## lavaan 0.6-9 ended normally after 25 iterations
##
##   Estimator                                         ML
##   Optimization method                           NLMINB
##   Number of model parameters                        27
##
##   Number of observations                           500
##
## Model Test User Model:
##
##   Test statistic                                27.015
##   Degrees of freedom                                27
##   P-value (Chi-square)                           0.463
##
## Parameter Estimates:
##
##   Standard errors                             Standard
##   Information                                 Expected
##   Information saturated (h1) model          Structured
##
## Latent Variables:
##                    Estimate  Std.Err  z-value  P(>|z|)   Std.lv  Std.all
##   factor =~
##     x1                0.535    0.027   19.740    0.000    0.535    0.809
##     x2                0.485    0.057    8.455    0.000    0.485    0.398
##     x3                0.513    0.068    7.536    0.000    0.513    0.358
##     x4                0.435    0.062    7.069    0.000    0.435    0.337
##     x5                0.460    0.042   10.839    0.000    0.460    0.497
##     x6                0.456    0.031   14.511    0.000    0.456    0.636
##     x7                0.489    0.029   16.668    0.000    0.489    0.710
##     x8                0.461    0.056    8.288    0.000    0.461    0.391
##     x9                0.471    0.049    9.592    0.000    0.471    0.446
##
## Intercepts:
```

```
##                     Estimate  Std.Err  z-value  P(>|z|)   Std.lv  Std.all
##     factor            0.000                               0.000    0.000
##    .x1                0.017    0.030    0.568    0.570    0.017    0.025
##    .x2                0.048    0.054    0.890    0.373    0.048    0.040
##    .x3                0.025    0.064    0.393    0.694    0.025    0.018
##    .x4                0.042    0.058    0.734    0.463    0.042    0.033
##    .x5               -0.043    0.041   -1.048    0.295   -0.043   -0.047
##    .x6                0.019    0.032    0.585    0.558    0.019    0.026
##    .x7                0.016    0.031    0.531    0.595    0.016    0.024
##    .x8               -0.041    0.053   -0.774    0.439   -0.041   -0.035
##    .x9               -0.001    0.047   -0.030    0.976   -0.001   -0.001
##
## Variances:
##                     Estimate  Std.Err  z-value  P(>|z|)   Std.lv  Std.all
##     factor            1.000                               1.000    1.000
##    .x1                0.151    0.016    9.362    0.000    0.151    0.345
##    .x2                1.249    0.082   15.188    0.000    1.249    0.842
##    .x3                1.790    0.117   15.326    0.000    1.790    0.872
##    .x4                1.479    0.096   15.388    0.000    1.479    0.886
##    .x5                0.643    0.044   14.714    0.000    0.643    0.753
##    .x6                0.307    0.023   13.499    0.000    0.307    0.595
##    .x7                0.234    0.019   12.272    0.000    0.234    0.495
##    .x8                1.180    0.078   15.215    0.000    1.180    0.847
##    .x9                0.892    0.060   14.985    0.000    0.892    0.801
dat02 = dat02 %>%
  mutate(factor.score = lavPredict(fit02))

low.var.sum02 = lm(scale(out) ~ scale(sum), data=dat02)
low.var.factor02 = lm(scale(out) ~ scale(factor.score), data=dat02)
```

**High Standardized Factor Loading Variance**

| | **Outcome ~ Sum Score** | | | **Outcome ~ Factor Score** | | |
|---|---|---|---|---|---|---|
| *Predictors* | *Estimates* | *SE* | *P-Value* | *Estimates* | *SE* | *P-Value* |
| Intercept | 0.000 | 0.036 | 1.000 | 0.000 | 0.034 | 1.000 |
| Sum Score | 0.581 | 0.036 | **<0.001** | | | |
| Factor Score | | | | 0.643 | 0.034 | **<0.001** |
| Observations | 500 | | | 500 | | |
| $R^2$ / $R^2$ adjusted | 0.337 / 0.336 | | | 0.414 / 0.413 | | |
